# Sensory neurons inhibit invadopodia and metastasis via direct CGRP-RAMP1-cAMP signaling to cancer cells

**DOI:** 10.64898/2026.04.17.719233

**Authors:** Ines Velazquez-Quesada, Elizaveta Belova, Afrooz Jarrah, Maria Carolina Cesar Mariano, Yasmine Dahleh, Maíra de Assis Lima, Debora Barbosa Vendramini Costa, Ralph Francescone, Edna Cukierman, Louis Hodgson, Bojana Gligorijevic

## Abstract

Breast cancer is globally the most common cancer among women. Although the five-year survival rate exceeds 80% for patients with localized disease, it drops to approximately 30% once metastasis occurs, underscoring the urgent need to define mechanisms that drive metastatic progression. Breast is a highly innervated organ and most of its innervation is sensory. However, whether sensory neurons can directly impact breast cancer cells remains an understudied topic.

Here, we show that mammary tumors have increased CGRP⁺ sensory innervation. Using our novel microfluidic Device for Cancer cell–Axon Interaction Testing (DACIT), we demonstrate that the presence of axons strongly inhibits ECM-degrading ability of cancer cells. The sensory neuron secretome suppresses assembly and function of invadopodia, which are cancer cell protrusions controlling ECM degradation, and essential for intravasation and metastasis. We identify calcitonin gene-related peptide (CGRP) as the key component of the sensory neuron secretome responsible for the inhibitory effect.

CGRP signaling occurs through the CRLR/RAMP1 receptor complex expressed by breast cancer cells, inducing a rapid increase in intracellular cAMP levels in breast cancer cells, followed by an increase in RhoC activity and suppression of invadopodia and ECM degradation. Loss of RAMP1 function enhances 3D spheroid invasion, cancer cell motility *in vivo* and significantly increases the number and the size of lung metastatic foci. Consistently, *in silico* analyses of both mouse and human RNASeq data point to a link between increasingly invasive subtypes with a gradual decrease in expression of RAMP1 and CRLR. To validate *in silico* findings, we compare RAMP1 expression in the patient breast tumors with adjacent normal tissues, confirming the invasive breast tumors have reduced levels of RAMP1.

Together, our findings identify a protective role for the paracrine CGRP signaling in limiting breast cancer invasion and metastasis. We also demonstrate how cancer cells circumvent CGRP inhibition by suppressing RAMP1 expression, highlighting CGRP-RAMP1-cAMP axis as a potential therapeutic target in breast cancer.

## Introduction

Metastatic breast cancer occurs in approximately 30% of breast cancer patients, yet it accounts for 70% of breast cancer-related deaths^1^. Thus, identifying mechanisms that influence cancer metastasis are of critical importance.

Peripheral innervation was recently reported abundant in melanoma, gastric, pancreatic, and breast cancers^1^. In human breast tumors, nerve fiber density and fascicle (axon bundle) size are higher compared to normal breast tissues and correlate with shorter disease-free survival ^2^, with innervation increasing further in invasive carcinomas compared to carcinomas *in situ* ^3^. Collectively, these data suggest that breast tumor innervation intensifies over the course of tumor progression ^2^.

Early studies of tumor-associated sensory neurons have revealed that their impact on tumor cell behaviors is cancer-specific, neuron subtype-specific and complex, sometimes yielding contradictory conclusions. In breast cancer specifically, Erin et al. have shown that surgical and chemical denervation of sensory neurons in syngeneic mammary tumors has no effect on primary tumor growth but significantly increases metastasis to the lungs and heart ^4,5^. Other studies have indicated that sensory neurons may promote breast tumor progression. For example, Padmanaban et al. showed that sensory neuron denervation in breast reduced tumor growth and lung metastases ^6^, whereas Le TT et al. demonstrated that contact with sensory neurons enhances cancer cell adhesion and migration via axon guidance molecule plexinB ^7^.

These apparently contradictory effects could be explained by the complexity introduced by two main factors. One, *in vivo* tumor denervation studies nonspecifically impact the whole organ, including host and tumor cells. As there is an intimate coordination between sensory neurons and host cell activities, such as immune cells^8–11^, results of denervation present the confounding impact on all cells in the tumor. Two, due to the general lack of cell lines properly modeling sensory neurons, *in vitro* studies are generally based on the use of primary dorsal root ganglion (DRG) and cancer cell co-cultures, which include many non-neuronal cell types (Schwann, immune, satellite cells) known to influence cancer cell behavior^12–14^.

Here, we focused on isolating interactions between sensory neurons and breast cancer cells and their impact on breast cancer invasion and metastasis. We used high-resolution microscopy of cleared tissues to show that CGRP+ sensory axons surround tumor vasculature. To interrogate the role of sensory neurons in regulating breast cancer cell invasion, we employed our Device for Cancer cell–Axon Interactions Testing (DACIT)^15,16^, an anatomically relevant microfluidic platform that physically separates sensory neuron soma and other DRG cell types from extending axons and cancer cells. While the compartmentalization maintains soma within the microenvironment important to their native functions, it eliminates confounding contributions from non-neuronal cells. Using DACIT, we show that the CGRP released by sensory axons suppresses invadopodia and ECM degradation. Mechanistically, CGRP elevates intracellular cAMP and RhoC activity in cancer cells, which leads to inhibition of invadopodia. Loss-of-function assays of CGRP receptor component RAMP1 show increase in 3D spheroid invasion *in vitro*, as well as increased cell movement and lung metastases *in vivo*. Finally, mouse and human *in silico* RNASeq analysis, validated by patient tissue microarrays (TMA), reveals that the CGRP receptor expression is decreased in aggressive tumors.

Together, these findings identify a previously unrecognized protective function of CGRP⁺ sensory axons on invadopodia and metastasis and suggest that attenuation of the CGRP/RAMP1 signaling axis may contribute to metastatic progression in patients with high invadopodia frequencies.

## Material and methods

### Ethical statement for animal experiments

This study was performed following the NIH regulations. The animal protocol was approved by the Temple University Ethical Committee (IACUC, protocol number 5072).

### Mouse models

Mice were purchased from Jackson Laboratories (Balb/c, stock# 000651; C57BL/6J, stock# 000664; FVB/NJ, stock# 001800; FVB/N-Tg 634Mul/J, stock# 002374-PyMT) or Charles Rivers Laboratories (SCID, CB17/Icr-*Prkdc^scid^*/IcrIcoCrl, Strain code: 236).

#### Xenografts

Cell suspension containing 10^6^ cells diluted in 100 µl PBS/Matrigel (1:1) was injected into the fourth mammary gland of 6-10-week-old female SCID mice (Charles River). Tumor growth was monitored by palpation weekly and upon palpation, and measured by calipers. Animals were sacrificed when the tumors reached 10 mm in diameter.

#### Intravital imaging

Mice with 8 mm mammary tumors (30-50 days post-injection) were surgically prepared for intravital imaging as previously described ^17,18^. Briefly, mice were anesthetized with 5%, followed by 2% of isoflurane, and the tumor was exposed by skin flap surgery. Mice were kept at 35°C in the *InVivo* environmental chamber. 4D stacks (100 µm in z; 5 min interval for 1 hour) were collected with a confocal-multiphoton microscope (Olympus FV1200MPE), with a 30X objective (UPLSAPO 30×, NA 1.05, Olympus). Areas were selected based on the presence of flowing vessels and high density of fluorescent cancer cells.

One-hour time-lapse videos were analyzed with FIJI. Single cells (shControl: 25 cells, sh1-RAMP1: 27 cells) of one field of view per model were manually selected at each time point (every 5 min), the centroid position determined and average cell speed calculated.

### Tissue collection and analysis

Tumors and lungs were collected and soaked in 4% paraformaldehyde (Alfa Aesar, Cat# 43368-9M) /PBS for 72 h. Then, the tissue was washed with PBS and maintained in ethanol 70% until use. Tumor weight was determined with a balance.

#### Lung metastasis quantification

Fixed lungs were imaged with Olympus FV3000 confocal microscope at 10X. Lungs were separated into lobes, nuclei were stained with DraQ5 (ThermoScientific, Cat#62251) and 3-5 lobes per mouse were fully scanned, collecting 3D stacks of Dendra2-fluorescent signal. Each field of view was combined into a maximum projection image, thresholded, and quantified for number and area of metastatic foci.

### DRG collection and dissociation

DRG dissection and dissociation from 6-12-week-old female Balb/c mice were performed as described previously ^15,19^. Briefly, animals were euthanized, the spinal cord was removed, and up to 30 cervical, thoracic, and lumbar DRGs were collected per mouse. DRGs were digested with 5 mg/ml Collagenase P (Sigma-Aldrich, Cat#11213865001) for 45 min at 37°C, followed by 5 min digestion with Trypsin/EDTA (Thermo Fisher Scientific, Cat# 25300120) at 37°C. Trypsin was inactivated with horse serum (Atlanta, Cat#S12150) in Neurobasal medium (Thermo Fisher Scientific, Cat# 21103049) and the cells were disaggregated using polished pipets. Cell suspension was filtered through a 40 µm cell strainer (Corning, Cat# 352340) to remove cell clusters and through a layer of 10% bovine serum albumin (BSA), to reduce the number of non-neuronal cells. Finally, neurons were concentrated by centrifugation (5 min, 0.1 x *g* at 20-22°C) and resuspended in complete neuron media: Neurobasal medium (Thermo Fisher, Cat# 21103049) supplemented with B27 (Thermo Fisher Scientific, Cat# 17504044), 10 U/ml penicillin, and 10 µg/ml streptomycin (Gibco, Cat#15140122) and Amphotericin B (Thermo Fisher Scientific, Cat# 15290026); and plated on coated DACITs, coverslips or plates.

### DRG-conditioned medium (DRG-CM)

Five thousand DRG cells were plated on 96-well plates coated with Matrigel (1:10) and cultured in neuron complete media (Neurobasal media supplemented with B27, pen/strep and Amphotericin B). At 1 day *in vitro* (DIV), medium was refreshed with 50 ng/ml of murine nerve growth factor (NGF, ThermoFisher Scientific, Cat#13257-019). At 5 DIV, the neural media was replaced with fresh cancer cell media (DMEM supplemented with 10% FBS and pen/strep or fresh complete neuron media with 1:500 Proteinase inhibitor Cocktail, (PIC; Sigma-Aldrich, Cat#P1860). Medium on coated wells without cells was used as controls. After 48 h, control and DRG-CM were collected for further analyses.

To test the effect of DRG-CM on the invadopodia degradation, freshly recovered control or DRG-CM were centrifuged 5 min at 5 000 x *g* at 20-22°C, diluted 2:1 with fresh medium and added to the cancer cells plated on gelatin.

### Cell lines and cell culture

MDA-MB-231 and BT549 cells were obtained from the Fox Chase Cancer Center cell culture facility (Philadelphia, PA, US) following authenticity (Short Tandem Repeat) and mycoplasma testing. Cells were maintained in DMEM (MDA-MB-231) or RPMI (BT549) medium supplemented with 10% Fetal Bovine Serum (FBS, Atlanta Biologicals), 10 U/ml penicillin, and 10 µg/ml streptomycin (Gibco, Cat#15140122) and incubated at 37°C at 5% CO_2_. MDA-MB-231 expressing Dendra2, MDA-MB-231-D2, were described previously^18^.

MDA-MB-231 stable knockdown sublines were generated using lentiviral particles from the Sigma-Aldrich MISSION library, targeting RAMP1 (sh1: TRCN0000014210; sh2: TRCN0000014212) and its non-targeting control (SHC202V), following the provider’s instructions. Transduced cells were selected for 7 days and further maintained with 1 µg/ml Puromycin (Cat# ICN10055210, MP Biomedicals).

### Antibodies

| Target | Source | Cat# | RRID | Dilution |
| --- | --- | --- | --- | --- |
| CGRP | Novus Biologicals | NB036540 | AB_3430321 | 1:100 |
| CGRP | Sigma-Aldrich | C8198 | AB_259091 | 1:400 |
| CK8.18 | Progen Biotechnik | GP11 | AB_1541064 | 1:200 |
| Cortactin | Abcam | ab81208 | AB_1640383 | 1:200 |
| Endomucin | Santa Cruz biotechnology | sc-65495 | B_2100037 | 1:400 |
| GAPDH | Santa Cruz biotechnology | Sc-47724 | AB_627678 | 1:1000 |
| PGP9.5 | BioSB | BSB 2115 | AB_3720319 | 1:100 |
| PGP9.5 | Abcam | ab15503 | AB_301912 | 1:200 |
| RAMP1 | Millipore Sigma | MABS1904 | AB_3741012 | 1:100 |
| RAMP1 | Abcam | ab203282 | AB_2924360 | 1:1000 |
| Tubulin βIII | BioLegend | 801202 | AB_2313773 | 1:1000 |
| β-actin | Santa Cruz biotechnology | sc-47778 | AB_2714189 | 1:1000 |
| CRLR | Bioss | bs-1860R | AB_10855106 | 1:500 |

### FRET biosensor for RhoC GTPase and ratiometric imaging

The optimized Förster Resonance Energy Transfer (FRET)-based biosensor for RhoC GTPase was derived from the previously published system ^20^. Briefly, the FRET donor mCerulean and the internal linker sections were modified using the synonymous codons to prevent random homologous recombination and biosensor truncation when expressed in living cells ^21^. Additionally, the FRET acceptor was modified from the original mVenus fluorescent protein to a circularly permutated (cp173) version of mVenus ^22^, improving the total dynamic range of FRET modulation. The biosensor expression cassette was inserted into NcoI and XhoI restriction sites of the pTriEX-4 backbone (Sigma, Cat#70824-3) and was transfected into MDA-MB-231 cells which were plated overnight at 2x10^5^ cells per 25 mm gelatin coated^23^ #1.5 round coverslips in 6-well cell culture plates, using Lipofectamine 2000 (Life Technologies, Cat#11668019) following the manufacturer’s protocols. Twenty-four hours following the transfection, cells were stimulated in the presence of 10% FBS with 40 μM forskolin (Sigma-Aldrich, Cat# F3917) in DMSO made at 40mM stock concentration, at 37 °C in 5% CO_2_ incubator, and fixed at the indicated time intervals using 1 % formaldehyde in PBS for 15 minutes. Following the fixation, cells were washed 3 times in PBS and immediately imaged for biosensor fluorescence.

Widefield FRET imaging was carried out using a custom Olympus IX83-ZDC2 microscope (Evident Scientific, Waltham, MA, USA) optimized for ratiometric biosensor analysis ^24^. Image acquisition was controlled using Visiview version 7.0.0.9 (Visitron Systems GmBH, Puchheim, Germany). Images were acquired through a 60 × 1.45 NA oil-immersion DIC objective lens under 2 × 2 camera binning. Simultaneous acquisition of cyan and yellow fluorescence emissions was achieved using two synchronized PrimeBSI-Express sCMOS cameras (Photometrics, Tucson, AZ, USA) mounted on a beamsplitter (Cairn Research Ltd., Faversham, Kent, UK) attached to the left-side emission port of the microscope. The following dichoric mirrors and band-pass filters were used: ZT440/514/561/640rpc-UF1, T560LPXRXT-UF2, and T505LPXR-UF2 dichroic mirrors, ET436/20X and ET480/40M for cyan fluorescent protein excitation and emission, and ET535/30M for yellow fluorescent protein FRET emission (Chroma Technology, Rockingham, VT, USA). Lumencor Sola-U-N solid-state illuminator was used for widefield illumination (Lumencor, Beaverton, OR, USA). During excitation of the field of view through the ET436/20X filter for the FRET donor excitation, the field illumination was attenuated through a neutral density filter ND 1.3 (Chroma Technology, Rockingham, VT, USA) and cyan and FRET emissions were acquired simultaneously using 100 ms exposure time for both cameras, which resulted in approximately 50-75% fill level of the 16-bit intensity range of the cameras. All image channels were registered prior to ratiometric calculations using pixel-by-pixel alignment based on *a priori* calibration and a non-linear coordinate transformation ^25^. Image processing was performed using MetaMorph version 7.10.5.476 (Molecular Devices, San Jose, CA, USA) and MATLAB version 2011a (MathWorks, Natick, MA, USA) and included flat-field correction, background and camera noise subtraction, threshold masking, ratiometric calculations, and spatial registration, as previously described^25^.

### Invadopodia-mediated gelatin degradation assay

#### On 48-well plates

Glass bottom 48-well plates (MatTek, Cat#P48G-1.5-6-F) were cleaned with 1N HCl for 10 min and coated with 50 µg/ml Poly-L-Lysine (Sigma, Cat#P8920) for 20 min, followed with previously prepared 0.2% fluorescent gelatin (Sigma, Cat#G2500) for 20 min. Excess gelatin was washed 3 times with PBS and gelatin was crosslinked with 0.2% glutaraldehyde for 20 min on ice. Glutaraldehyde was quenched with 5 mg/ml sodium borohydride for 15 min at RT. Excess was washed out with PBS. For storage, penicillin-streptomycin (Pen/Strep) in PBS was added and plates were kept at 4°C protected from light until use.

On the day of the experiment, 25,000 cells/well were plated and left to attach for 1 h at 37°C, 5% CO_2_. When mentioned, cells were then treated with 5 nM – 50 µM CGRP (Anaspec, Cat#AS-20681), 5 µM KP10 (Anaspec, Cat# AS-64240), 5 µM Substance P (Anaspec, Cat#AS-24279), 5 µM Neuropeptide Y (Tocris, Cat#1153), 40 µM Forskolin (Sigma-Aldrich, Cat# F3917), 100 µM 3-isobutyl-1-methylxanthine (IBMX; Sigma-Aldrich, Cat#I7018), or control or DRG-conditioned media (DRG-CM). Restored compounds were aliquoted and storage at -20 or -80°C, and freeze-thaw cycles were minimized. After 24 h, cells were fixed with 4% paraformaldehyde ( Alpha Aesar, Cat# 43368-9M) for 10 minutes and plates were imaged.

#### In DACIT

We prepared DACIT as previously described ^15,16^. Briefly, a 10:1 ratio of Sylgard 184 elastomer base and curing agent (Electron Microscopy Sciences) was mixed and poured into a custom-fabricated SU-8 DACIT master, which was then baked overnight at 65°C. The next day, the cured polydimethylsiloxane (PDMS) DACITs were cut, plasma-treated and sealed by plasma-treated glass coverslips (24 mm squares, #1405-10, Globe Scientific Inc). Assembled DACITs were sterilized with 70% ethanol and 15 min of UV light exposure. All the procedures were done in the laminar hood at RT unless otherwise stated. The neuronal compartment was coated with 50 µg/ml Poly-L-Lysine overnight, followed by 1:10 Matrigel in PBS (Corning, Cat# 356234), for 30 min at 37°C. The axonal compartment was coated with fluorescent gelatin and topped with laminin (Sigma, Cat#L2020), as previously reported ^15^. Briefly, procedure described for well plates was followed, but the glutaraldehyde was quenched for 15 min using 50 mM Glycine (Alfa-Aesar, Cat# J62407-22). Next, the axonal compartment was once again treated with 50 µg/ml of Poly-L-Lysine for 20 min and topped with a layer of 5 µg/ml laminin for 20 min.

Between 10-20,000 primary DRG neurons were plated on the neuronal compartment. The following day, the medium was refreshed, and 0.5 µg/ml of NGF (ThermoScientific Cat#13257019) was added to the axonal compartment only, maintaining the fluidic compartmentalization. At 3 DIV, the medium from the axonal compartment was removed, the channel was washed twice with HBSS, Ca^++^, Mg^++^ free (Thermo Fisher Scientific, Cat# 14175095), and 7,500 MDA-MB-231 cells were plated, followed by addition of cancer cell media. At 6 DIV, cells were fixed by 4% paraformaldehyde for 10 minutes, and images of cells and gelatin layers were collected by confocal microscope.

Images of invadopodia, nuclei and fluorescent gelatin layer were acquired using laser scanning confocal microscope Olympus FV1200 equipped with a 60X oil immersion objective (UPLSAPO60XS, 1.35 NA, Olympus), or Nikon Eclipse Ti2 microscope with a 60X oil objective (Plan Apo 60X/1.4 oil, Nikon). Z-stacks were captured with 0.25 µm step sizes. Each experiment was done in biological triplicate and technical triplicate (48-well plate) or duplicate (DACIT), collecting 3D stacks of 10+ fields-of-view (FOV) per condition.

Cells with invadopodia precursors were identified as cells with puncta with colocalization of cortactin and actin, while cells with invadopodia were identified as actin-positive puncta colocalizing on degraded gelatin. Gelatin channel was analyzed using a custom macro in Fiji, first compiling the Z-stack slices into a Min Intensity projection, and then thresholding the signal. Degradation puncta were identified using the Particle Analysis tool. To account for differences in cell density across the fields of view, the total degradation area was normalized by dividing it by the number of cells in the same field, with the number of cells determined based on the nuclei staining or brightfield.

### Cell viability

Crystal violet staining was employed to assess the proliferation of MDA-MB-231 sublines. Briefly, cells were seeded in 96-well plates at 10,000 cells/ml. At designated time points, cells were washed once with cold PBS, fixed with 100% cold methanol, and stained with a 0.5% crystal violet solution (Acros Organics, Cat# 405830250) prepared in 25% methanol. After a 10-minute incubation, excess dye was removed, and wells were rinsed rigorously with tap water. Following air-drying, the bound dye was solubilized in 100% methanol, and absorbance was measured at 570 nm using an Infinite M200 Pro Tecan plate reader. The cell density at 12 hours post-seeding was designated as day 0.

### Time-lapse recording of G-Flamp1 in cancer cells

The pCAG-G-Flamp1 plasmid (donated from Jun Chu, Addgene plasmid #188567; http://n2t.net/addgene:188567; RRID:Addgene 188567)^26^ was transiently transfected into the MDA-MB-231 cells using Lipofectamine 3000 (Thermo Fisher Scientific, Cat#L3000008), following the recommendations of the provider. After 48 h, the media was replaced with fresh live imaging solution (Gibco, A59688DJ) and imaged. Time-lapse images were collected every 15 sec for 15 minutes (60 time-points) on the Nikon Eclipse Ti2 microscope, using the 10X objective, adding treatments at minute one.

Live images were analyzed with Fiji (ImageJ, v2.16.0), and data plotted in GraphPad Prism (v10.6.1). Briefly, the average fluorescence intensity of each region of interest (ROI) was determined with Fiji for each time point, and the Δ*F*/*F*_0_ was calculated using the formula:

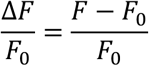

In which, ***F*** is the fluorescence intensity at each time point, and ***F*_0_** is the baseline fluorescence intensity defined as the average fluorescence of the first 6 timepoints.

### 3D Spheroid Formation and Invasion

Spheroids were generated as previously described ^27^. MDA-MB-231 Ctrl and shRAMP1 cells were trypsinized and resuspended at 1 × 10⁴ cells/ml in spheroid formation medium (complete DMEM/F12 supplemented with 5% horse serum, 0.5 µg/ml hydrocortisone, 20 ng/ml human EGF, 10 µg/ml insulin, 100 ng/ml cholera toxin, and 1% penicillin/streptomycin). Two hundred microliters of medium containing cells (2,000 cells per well) were seeded into non-adherent round-bottom 96-well plates (Costar, Cat#3788), centrifuged at 1000 rpm for 5 min at room temperature, and placed on an orbital shaker (∼60 rpm) for 2 h at 37°C. After 2 h, the spheroid formation medium was removed, and 200 µl of cold spheroid growth medium (complete DMEM/F12 supplemented with 0.25% methylcellulose and 1% Matrigel) was gently added to each well. Cells were incubated under static conditions for 48 h to allow spheroid maturation.

After 48 h, spheroids were collected into complete medium (high-glucose DMEM supplemented with 10% FBS and 1% penicillin/streptomycin) and embedded in 30 µl of a 3 mg/ml collagen I matrix as previously described ^28^. Briefly, high-concentration Collagen I (Corning, Cat#354249) was diluted on ice to 3 mg/ml final concentration using 10X PBS (Thermo Fisher Scientific, Cat#14200166) (added at 0.1x the final volume), 1 N NaOH (Thermo Fisher Scientific, SS266-1) (0.023 ml per ml of Collagen I) to neutralize the solution, and ice-cold complete medium to adjust the final volume. Thirty microliters of the collagen I mixture containing spheroids was added to the wells of the spheroid imaging device ^29^ and allowed to polymerize at 37°C for 30 min, after which complete medium was added to the dishes. Spheroids were allowed to invade the collagen matrix for 72 h, then fixed.

Spheroids were imaged on day 0 and day 3 using an Olympus FV3000 confocal microscope, acquiring z-stacks at 10 µm intervals. Images were quantified using Fiji. Briefly, images were maximum-intensity projected, and the spheroid area was determined using the wand tool on the Dendra channel. The area growth, defined as the spheroid area at day 3 divided by the average spheroid area at day 0, was normalized to the shCtrl median condition (relative area).

### Western blots

Cells were harvested in RIPA buffer (Teknova, Cat# R3792) supplemented with protease-(Cat#11836170001, Sigma-Aldrich), and phosphatase inhibitor cocktails (ThermoFisher Scientific, Cat#78420). Protein amounts were quantified with the BCA Protein Assay (ThermoFisher Scientific, Cat# 23227) and 30 µg of proteins were loaded into 12% pre-casted gels (Mini-PROTEAN TGX gels, Bio-Rad) and run at 100V until the proteins enter to the resolving gel and at 150 V until the dye front reaches the bottom of the gel. After running, proteins were transferred to a PVDF membrane (Sigma Aldrich, Cat# IPVH00005) using a semi-dry transfer system (Bio-Rad, Trans-Blot SD semi-dry transfer cell) at 20 V for 15-30 min. Nonspecific sites were blocked with 5% BSA in TBS-T (0.1% Tween 20 in Tris-buffered Saline) for 1 h at 20-22°C, incubated with primary antibodies diluted in 1.5% BSA in TBS-T overnight at 4°C. Membranes were washed with TBS-T and incubated with HRP-linked secondary antibodies for 1 h at 20-22°C. Proteins were visualized using chemiluminescence, using either WesternBright ECL HRP substrate (Advansta, Cat# K-12045-D20), or SuperSignal™ West Femto Maximum Sensitivity (Thermo Scientific, Cat# 34094) and visualized on a blot scanner (C-DiGit, LI-COR). Protein band intensities were quantified with ImageJ.

### Immunofluorescence

#### Cells

Plated cells were fixed with 4% paraformaldehyde for 10 min, washed and permeabilized for 5 min with Triton X-100 (0.1%; EMD Millipore, Cat# 94-001-00ML) in PBS. To block nonspecific sites, cells were incubated for 1 h with Intercept® Blocking Buffer (LICORbio; Cat#927-70001). Antibodies diluted in blocking solution were incubated with the cells overnight at 4°C. The next day, plates were washed with PBS and incubated for 1 h RT with the secondary antibody diluted in PBS. Plates were washed again with PBS and mounted with Fluoromount-G^TM^ Mounting Medium (ThermoFisher, Cat#00-4958-02).

#### Paraffin-embedded tissue

Paraffin-embedded tissue was sectioned at 5 µm and mounted onto glass slides. Prior to staining, sections were deparaffinized and subjected to antigen retrieval by incubation in R-UAR buffer (Electron Microscopy Sciences, Cat# 62719-10) for 20 min at 120 °C. After cooling, immunofluorescence staining was performed as previously described for cultured cells. Before mounting, slides were consecutively dehydrated and mounted using Fluoromount-G™ Mounting Medium.

### Immunohistochemistry

We performed immunohistochemistry on tissue as previously described^19^. Briefly, murine mammary glands and PyMT mammary tumors were collected and fixed in 4% paraformaldehyde for 48 h, dehydrated and embedded in paraffin. Immunohistochemistry staining was performed on 5 µm sections using VENTANA Discovery XT automated staining (Ventana Medical Systems) according to the manufacturer’s instructions. Briefly, slides were deparaffinised using EZ Prep solution (Ventana, Cat# 950−102) for 16 min at 72°C. Epitope retrieval was accomplished with CC1 solution (EDTA, pH 9.0. Cat# 950−224) at high temperature (95−100°C) for 32 min. Mouse primary antibody (Tubulin β III, 1:1000, Cat #801202. BioLegend) titered with a TBS antibody diluent into user-fillable dispensers for use on the automated stainer. Immune complex was detected using the Ventana OmniMap anti-mouse detection kit (Cat# 760-4310) and developed using the VENTANA ChromMap DAB detection kit (Cat# 760-159). Slides were counterstained with hematoxylin II (Cat# 790–2208) for 8 min, followed by Bluing reagent (Cat# 760–2037) for 4 min. The slides were then dehydrated with ethanol series, cleared in xylene and mounted.

### Tissue microarray

#### Slide preparation and immunohistochemistry

Immunohistochemical staining for RAMP1 (Receptor Activity-Modifying Protein 1) was performed on formalin-fixed, paraffin-embedded (FFPE) human breast tissue array section (TissueArrray.com, Cat# BC08118b).

Tissue sections were dried overnight at 50 °C prior to staining. Then, the slide was deparaffinized using EZ Prep solution (Ventana, Cat# 950−102) according to the manufacturer’s protocol. Heat-induced epitope retrieval was performed using Cell Conditioning 1 Buffer (CC1; Tris-EDTA buffer, pH 8.0) at 95°C for 32 min. Retrieval conditions were optimized during assay development to achieve optimal signal-to-noise ratio for RAMP1 detection. Primary antibodies (anti-RAMP1, 1:2000, Cat# ab156575) were added to the slide and incubated for 32 minutes at 37°C.

Detection was performed using a multimer-based horseradish peroxidase (HRP) system (Discovery OmniMap Anti-Rabbit HRP). Signal visualization was achieved using the Discovery ChromoMap DAB IHC Detection Kit, with 3,3′-diaminobenzidine (DAB) serving as the chromogenic substrate to produce a brown precipitate at antigen sites. Slide was subsequently counterstained with hematoxylin followed by application of a bluing reagent.

Appropriate positive control tissues known to express RAMP1 were included in each staining run. Negative controls were generated by substituting the primary antibody with species-matched non-immune IgG.

#### Slide Scanning

All stained sections were digitized as whole-slide images using a Hamamatsu NanoZoomer S60 slide scanner at 40× magnification under standard acquisition settings.

#### Image analysis using Visiopharm

Quantification of RAMP1 expression was performed using custom analysis applications developed in Visiopharm software (Visiopharm A/S, Denmark; version 2025.08, 64-bit). All applications were optimized to accommodate biomarker-specific staining characteristics and ensure reproducible measurements across samples.

#### Tumor detection

Tumor detection applications were developed to distinguish viable tumor regions from surrounding stromal and necrotic tissue. Quantitative analysis was restricted to automatically detected tumor areas to ensure consistent exclusion of non-tumor tissue.

The detection algorithms were customized to accommodate morphological variability across samples, enabling consistent tumor identification while minimizing inclusion of background or nonviable tissue.

#### H-Score application development

Following tumor region detection, color deconvolution was applied to separate hematoxylin and DAB channels. DAB-positive pixels were classified into high-, medium-, and low-intensity categories, while hematoxylin-positive pixels were detected for normalization.

Analysis parameters were optimized for stain intensity, cellular morphology, and background characteristics unique to each antibody. Threshold levels, color deconvolution settings, and tissue classification criteria were iteratively refined to achieve consistent and biologically meaningful segmentation across all samples.

Threshold values for both channels were optimized for RAMP1 to reflect biomarker-specific staining patterns. Areas corresponding to high-, medium-, and low-intensity DAB staining, along with hematoxylin-positive areas, were calculated.

Cytoplasmic pixel-based H-scores (Cyto H-scores) were computed using the formula:

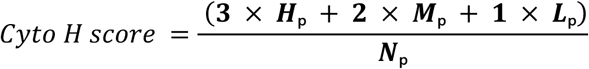

where H_p_, M_p_, and L_p_ represent areas of high-, medium-, and low-intensity DAB-positive pixels, respectively, and N_p_ represents the hematoxylin-positive area. Sequential analysis prioritized the DAB channel to ensure accurate cytoplasmic quantification while minimizing interference from nuclear staining.

### Tissue immunofluorescence and clearing

Tissue immunofluorescence followed by clearing was performed using SeeDB protocol, as previously described ^19,30^, with some modifications. Briefly, mammary glands (FVB mice) or mammary tumors (PyMT mice) were collected and fixed in 4% paraformaldehyde at 4°C for 72 hours, embedded in 4% low-melting agarose (Fisher Scientific, Cat# BP165), and sectioned at 500 µm using the TPI Vibratome series 1000 tissue sectioning system.

Tissue sections were then re-fixed and permeabilized for 3 hours at room temperature in PBS containing 5% (w/v) D-sucrose (Sigma, Cat# SLCG8436), 4% paraformaldehyde, and 0.5% Triton X-100. Excess fixative was removed by washing with PBS containing 0.1% (w/v) Tween-20 (Sigma, Cat# P7949). To block nonspecific sites, tissues were incubated for 48 hours at RT under gentle shaking (UNICO Test Tube Rocker) in PBS-GT (PBS supplemented with 0.2% gelatin and 0.5% Triton X-100). Sections were incubated with primary antibodies diluted in PBS-GT for 4 days, using intermittent centrifugation: 600 × *g* (Fisher Scientific accuSpin Micro 17) during the day, and gentle rocking (Fisher Scientific Incubating Mini Shaker) overnight. Excess primary antibodies were removed with three PBS-GT washes. Samples were then protected from light and incubated with secondary antibodies diluted in PBS-GT for 2 days under gentle rocking.

For refractive index matching, tissue sections were incubated in PBS with 0.5% (w/v) α-thioglycerol (Sigma-Aldrich, Cat# 88640) and increasing concentrations of fructose (20–115% w/v; Sigma-Aldrich, Cat# F0127): 8–16 hours at 20%, and 8–12 hours each at 40%, 60%, and 80%. Finally, tissues were incubated for 24 h at 100% and 115% fructose, under gentle rocking. Samples were stored in 115% fructose solution until imaging.

Cleared tissue was imaged using a confocal-multiphoton microscope (Olympus FV1200MPE), and 3D images were visualized using Imaris software (Imaris X64, v8.3.10).

### ELISA

To measure the concentration of CGRP in media from cancer cells or neurons, fresh cancer cell media with 1:500 Proteinase inhibitor Cocktail, (PIC; Sigma-Aldrich, Cat#P1860) was added to 5 DIV cells. After 48 h, DRG-CM was collected and the CGRP was quantified by ELISA (Cayman Chemical Company, Cat# 589001), following the instructions of the manufacturer.

### Single-cell RNA-seq analysis

We used the publicly available single-cell transcriptome datasets from Gene Expression Omnibus (GEO) **GSE221528** ^31^ and from **GSE75688** ^32^. Single-cell RNA-sequencing data were processed and analyzed in Python using Scanpy (v1.9.x). Raw expression matrices were imported into an AnnData framework, cell types were identified by unsupervised Leiden clustering on highly variable genes followed by annotation using canonical marker genes, and individual cells were assigned to experimental groups based on tumor collection time points: Day 11, Day 14, Day 18, and Day 24 for mouse samples, and on mammary tumor molecular subtypes: ER+ (Luminal), ER+HER2+ (Luminal B-like), HER2+, TNBC, LN metastasis ER+, HER2, LN metastasis TNBC for human samples. Cells lacking clear condition labels were excluded from further analysis.

Expression patterns of CGRP receptor–associated genes (RAMP1, CALCRL) were examined across tumor time points within each defined cell population. Raw counts from the mouse transcriptome (scRNAseq, GSE221528) were normalized to 10,000 counts per cell based on library size and subsequently log1p transformed prior to analysis. Pre-processed transcripts per million (TPM) values obtained from the transcriptome from breast cancer patients (scRNAseq, GSE75688) were transformed using log2(TPM+1).

For dot plot visualization, expression values were scaled per cell type using min-max normalization. The mean log-normalized expression was calculated across all cells pooled together within each cell type and condition group. The minimum mean expression value across all conditions within each cell type was set to 0 and the maximum to 1, applied independently per cell type. Colors therefore reflect relative expression dynamics within each cell type and cannot be compared across cell types or across plots. Dot size represents the fraction of cells with detectable expression (> 0), scaled to the range present in each individual plot. Dots with fewer than 1% of cells expressing are shown as empty circles. Relative intensities were determined with the formula:

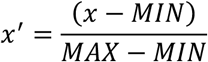

### Bulk RNAseq data analysis

Expression of *RAMP1* and *CALCRL* on mammary tumors major subtypes: Normal (n=114), Luminal (n=566), HER2+ (n=37), triple negative breast cancer (TNBC, n=116) was determined using RNAseq dataset from the University of Alabama at Birmingham CANcer data analysis Portal (UALCAN) ^33,34^.

### Quantification and statistical analysis

Images were analyzed with Fiji (ImageJ, v2.16.0) ^35^. Statistical analysis and graphs were made using GraphPad Prism (v10.6.1). Unless stated otherwise, all tests were performed using unpaired and two-sided criteria. Data are represented as violin plots. Statistical significance was defined as \**p* < 0.05, \*\**p* < 0.01, \*\*\**p* < 0.001, \*\*\*\**p* < 0.0001. Additional information can be found in the figure legend. All data are available upon request.

Cell segmentation was performed using the cyto model of Cellpose ^36^, (version 2.2.1; python version: 3.11.1). Segmented files were uploaded in Fiji to determine the number and shape parameters of segmented cells.

Representative images were selected from samples with measurements closest to the median (or mean) of the quantified dataset. Figure 4E was created with BioRender.

## Results

### Mammary tumors have larger fascicles and increased sensory innervation compared to mammary glands

To examine the innervation in mammary tumors, we used the PyMT-MMTV transgenic mouse model ^37,38^, which spontaneously develops multifocal invasive tumors across all mammary glands and in which the tumor microenvironment co-evolves with cancer progression^38^. Using the general neuronal marker Tubulin β3, we observed fascicles in the core and at the periphery of the tumor (**Figure 1A**). Quantification recapitulated clinical observations in breast tumors, showing a significant increase in fascicle size in mammary tumors compared to mammary glands (**Figure 1B**)^2,39^. This suggests that murine mammary tumors recruit additional axons to pre-existing mammary fascicles.

**Figure 1.**
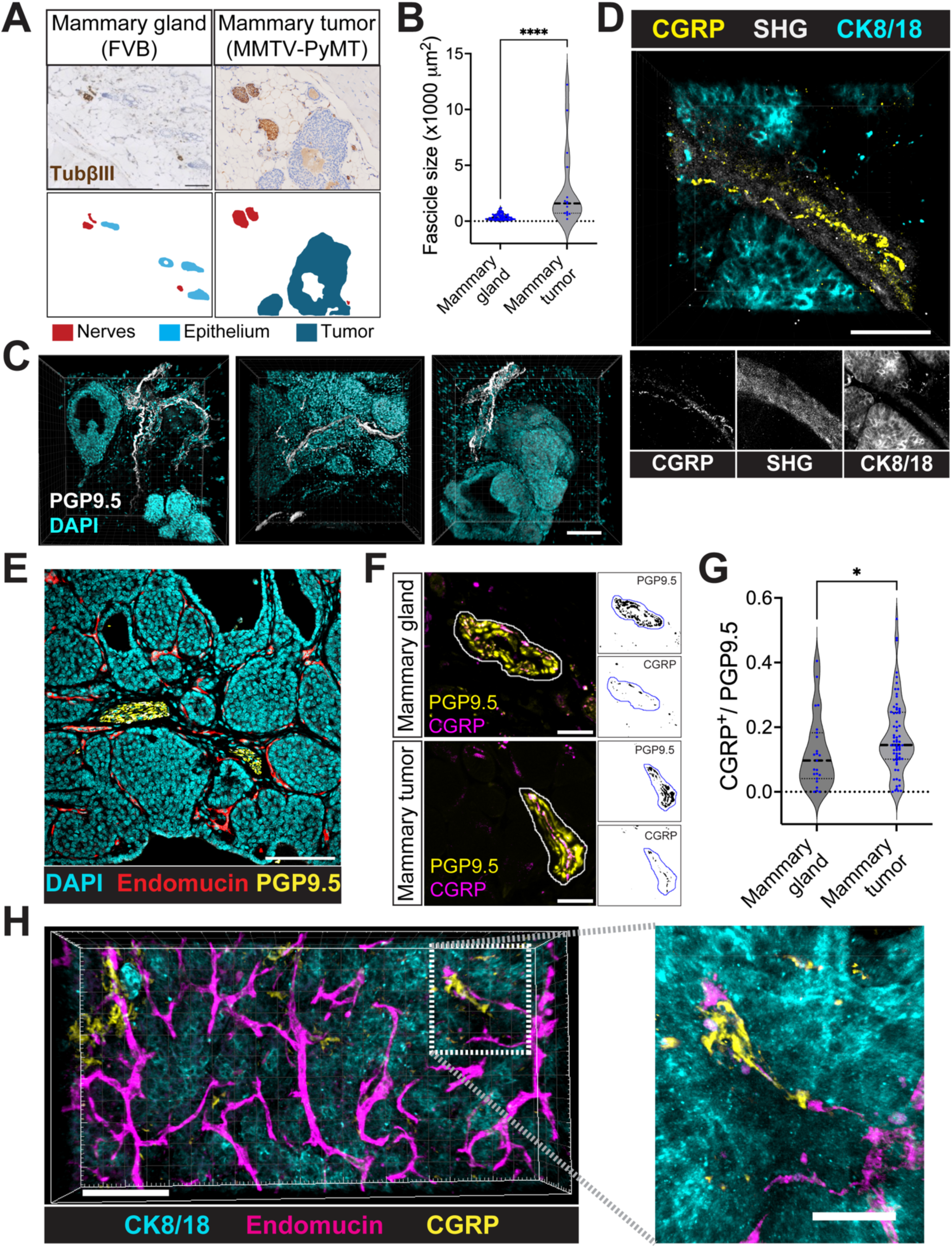
Overall innervation and sensory innervation are increased in the PyMT-MMTV mammary carcinoma. **A**. Immunohistochemistry (top) and scheme (bottom) showing fascicles (Tubβ3+, brown) on murine mammary gland (left) and PyMT mammary tumor (right). **B**. Graph showing quantification (median and quartiles) of the fascicle size on normal mammary gland (FVB; One mouse, 5 slides, n=60) and murine mammary tumors (PyMT; two mice, 2 slides, n=15). **C**. Immunofluorescence followed by clearing of the PyMT mammary tumor showing axons (PGP9.5+, white) and cell nuclei (DAPI, cyan). Scale bar = 100 µm. **D**. Immunofluorescence and clearing of PyMT mammary tumors showing CGRP (yellow), second harmonic-generation (SHG, white), and cytokeratin 8/18 (cyan). Scale bar = 50 µm. **E**. Immunofluorescence of cancer cells (CK8/18, cyan), vessels (Endomucin, red), and nerves (PGP9.5, yellow) on mammary PyMT tumor. Scale bar: 100 µm. **F**. Representative images of fascicles on the murine mammary gland (top) and PyMT mammary tumor (bottom) showing all (PGP9.5, yellow) and CGRP (CGRP, magenta) axons and mask. Scale bar: 25 µm. **G**. Fraction of CGRP+/PGP9.5+ signal (median and quartiles) on the murine mammary glands (FVB, N=2, n=23) or PyMT mammary tumors. (N=6, n=68). **H**. Immunofluorescence followed by clearing of PyMT mammary tumors. Cancer cells (CK8/18, cyan), vessels (Endomucin, magenta), and CGRP (yellow). Scale bar: 100 µm (left), 30 µm (right). Statistical differences were determined by the Mann-Whitney test. **** *p* < 0.0001; \**p* < 0.05.

To better resolve the spatial distribution of tumor innervation, we imaged cleared and immunolabeled tumor tissue in 3D at high-resolution. This approach revealed high axon frequency along the invasive edge (**Figure 1C**, left), within the tumor core (**Figure 1C**, middle), and close to structures resembling blood vessels (**Figure 1C**, right).

To specifically identify sensory neurons, we used the calcitonin gene-related peptide (CGRP), a neuropeptide expressed by peptidergic sensory neurons, that we showed innervate the mammary tumor^19^. We observed CGRP+ sensory neurons align with collagen fibers (**Figure 1D**). To determine the contribution of sensory neurons in the overall tumor innervation, we identified the fascicles within the tumor (**Figure 1E**) and quantified the proportion of CGRP+ fibers (**Figure 1F, magenta**) within PGP9.5+ fascicles (**Figure 1F, yellow**). Interestingly, the relative abundance of CGRP+ sensory fibers within nerve fascicles increased in mammary tumors compared with the mammary gland (**Figure 1F, 1G**), suggesting that breast tumors not only recruit additional axons to pre-existing mammary fascicles, but enrich fascicles with sensory axons. Subsequent 3D mapping of CGRP+ axons shows they are located in intimate proximity to blood vessels (endomucin+, **Figure 1H**), suggestive of a possible role in cancer cell intravasation^18^.

### Sensory neurons inhibit matrix degradation by cancer cells

To address the mechanisms of sensory axon interactions with cancer cells at the cellular level, we cultured adult mouse DRG neurons *in vitro*. It is known that in addition to neurons, ganglia contain a high proportion of macrophages, fibroblasts, as well as specialized satellite and Schwann cells^40^. Accordingly, while primary DRG cultures contain many neurons (PGP9.5^+^) that remain viable and extend axons in culture (**Supplementary Figure 1A)**, numerous non-neuronal cells (DAPI+/PGP9.5^-^) remain as part of the DRG primary culture supporting neuronal viability and function and comprising ∼97% of the total cells in culture (**Supplementary Figure 1B**).

To analyze direct axon-cancer cell interactions, while avoiding confounding signals from non-neuronal cells, we used DACIT (**Figure 2A**), a microfluidic device that physically separates DRG primary culture (left, yellow) from cancer cells (right, magenta), allowing only axons to reach the cancer cell compartment^15^ . We recently showed that DACIT mimics the axonogenesis from the DRG to the breast and preserves the proportion of sensory neuron subtypes which innervate mammary tumors^19^. Finally, through fluidic isolation, DACIT enriches axon-secreted factors within the axonal compartment ^15,16^.

**Figure 2.**
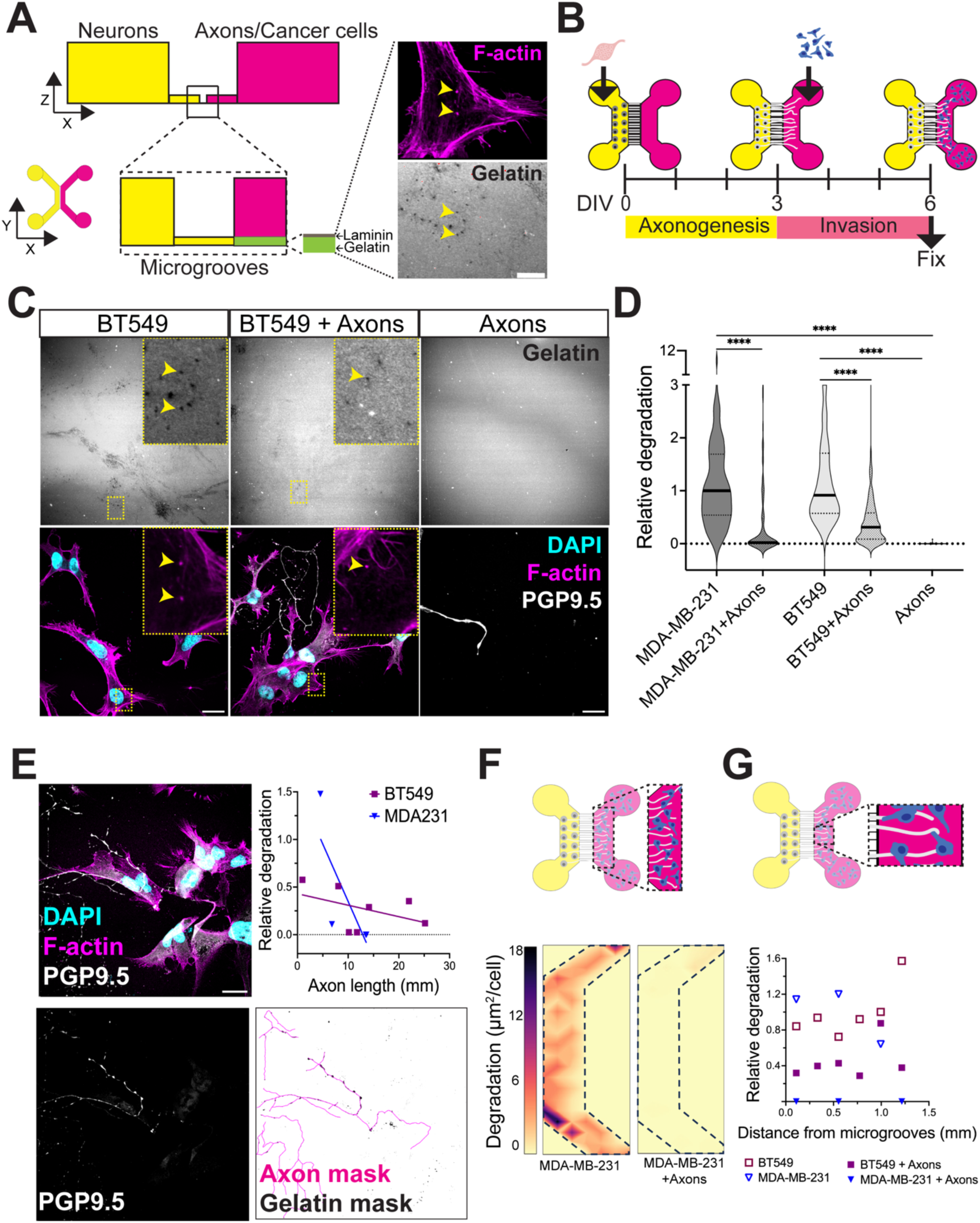
Presence of sensory axons inhibits the matrix degradation by cancer cells. **A**. Schematic of the Device for Axons-Cancer cell Interaction (DACIT) designed to study invadopodia formation on a gelatin degradation assay. The neuronal compartment (yellow) is connected to the axonal/cancer cell compartment (magenta) via microgrooves. The axonal/cancer cell compartment is coated with fluorescent gelatin and a laminin layer to assess cancer cell-mediated gelatin degradation. In the zoomed inset the arrows point F-actin-rich puncta (magenta, top) colocalizing with gelatin degradation spots (green, bottom). Scale bar = 25 µm. **B.** Experimental setup for the gelatin degradation assay in the DACIT. Primary cultures of murine dorsal root ganglia (DRG) neurons were seeded in the neuronal compartment. After 3 days *in vitro*, cancer cells were added to the axonal/cancer cell compartment. Two days later, cells were fixed, and gelatin degradation was quantified. **C.** Representative images from the axonal/cancer cell compartment, showing degraded gelatin (grays, top) and immunofluorescence labeling (bottom) of nuclei (cyan), F-actin (magenta), and axons (white). Scale bar = 25 µm. Yellow arrows point F-actin puncta colocalizing with degradation spots. **D.** Quantification of relative gelatin degradation by BT549 cells and MDA-MB-231 cells. Data represent three biological replicates (N=3), with two experimental replicates (n=2) and 10 fields-of-view analyzed per DACIT. Statistical significance was assessed using a two-tailed Mann Whitney test. *p* value < 0.05 are reported. **E.** Immunofluorescence showing BT549 cells (F-actin) with axons (PGP9.5) on the axonal compartment. Axons were traced using the PGP9.5 channel (bottom left) and the obtained mask (bottom right) was used to determine axon length on a degraded gelatin (top right). A Spearman test revealed a very strong (MDA-MB-231, *r* = -0.95) to strong (BT549, *r* = -0.41) negative relationship between axon length and gelatin degradation. **F.** Heatmap showing the gelatin degradation area (µm²/cell) across the DACIT. Left: cancer cells alone. Right: cancer cells in the presence of axons. **G.** Correlation of normalized gelatin degradation along the x-axis of the axonal compartment in the DACIT. The x-axis represents the distance from the microgrooves (mm), and the y-axis shows normalized gelatin degradation. Each point represents three biologicals with two experimental replicates. A Spearman correlation analysis revealed undetermined (MDA-MB-231 + Axons, *r* = ND – horizontal line) and weak (BT549 + Axons, *r* = 2) relationship between the distance from the microgrooves and the gelatin degradation.

To quantify invasion in cancer cells directly contacting sensory axons, we then performed the modified gelatin degradation assay in DACIT^41^, coating the axonal compartment with a layer of fluorescent gelatin and laminin to promote axonal attachment and detect degradation (**Figure 2A, B**). In this assay, gelatin degradation is detected as black spots on the gelatin channel that typically co-localize with F-actin-rich structures (**Figure 2A, yellow arrows**) and indicates the ability of cancer cells to form functional invadopodia.

Sensory axons did not affect cancer cell proliferation or their morphology (**Supplementary Figure 2).** However, the presence of axons significantly decreased the gelatin degradation in both BT549 (68% reduction) and MDA-MB-231 (72% reduction) breast cancer cells, while axons alone did not degrade matrix (**Figure 2C,D**).

We wondered whether local axon density was correlated with invadopodia inhibition. Pearson correlation analyses revealed that gelatin degradation negatively correlated with axon density (**Figure 2E**). However, invadopodia inhibition was homogenously distributed along the axonal compartment (**Figure 2F**), showing weak or no correlation with axon proximity (**Figure 2G**). This suggests that the inhibitory effect of sensory axons on matrix degradation was independent of direct contact and likely mediated by neuron-secreted factors.

### CGRP in the sensory neuron secretome inhibits invadopodia via CRLR/RAMP1

To test if the matrix degradation was inhibited by factors released by the sensory neurons (i.e. neurotransmitters), we plated MDA-MB-231 cancer cells on gelatin and cultured them either in control media, or in DRG-conditioned media (DRG-CM) defined as media conditioned by primary DRG cultures over 48 hours (**Figure 3A, 3B**). Indeed, DRG-CM reduced degradation by approximately 37% compared to control media (**Figure 3B**). We next quantified the number of invadopodia present on the membrane of cancer cells. Invadopodia assembly starts with punctate co-localization of F-actin and cortactin, and once precursors are assembled, they mature and mediate matrix degradation. Importantly, the sensory neuron secretome seemed to affect the early stages of invadopodia assembly, as the number of overlapping cortactin/F-actin spots (**Figure 3C, see line plot**) defined as invadopodia precursors was significantly decreased (**Figure 3D**). Together, these data suggest that the sensory secretome inhibits assembly of invadopodia, consequently resulting in reduction of invadopodia-mediated degradation.

**Figure 3.**
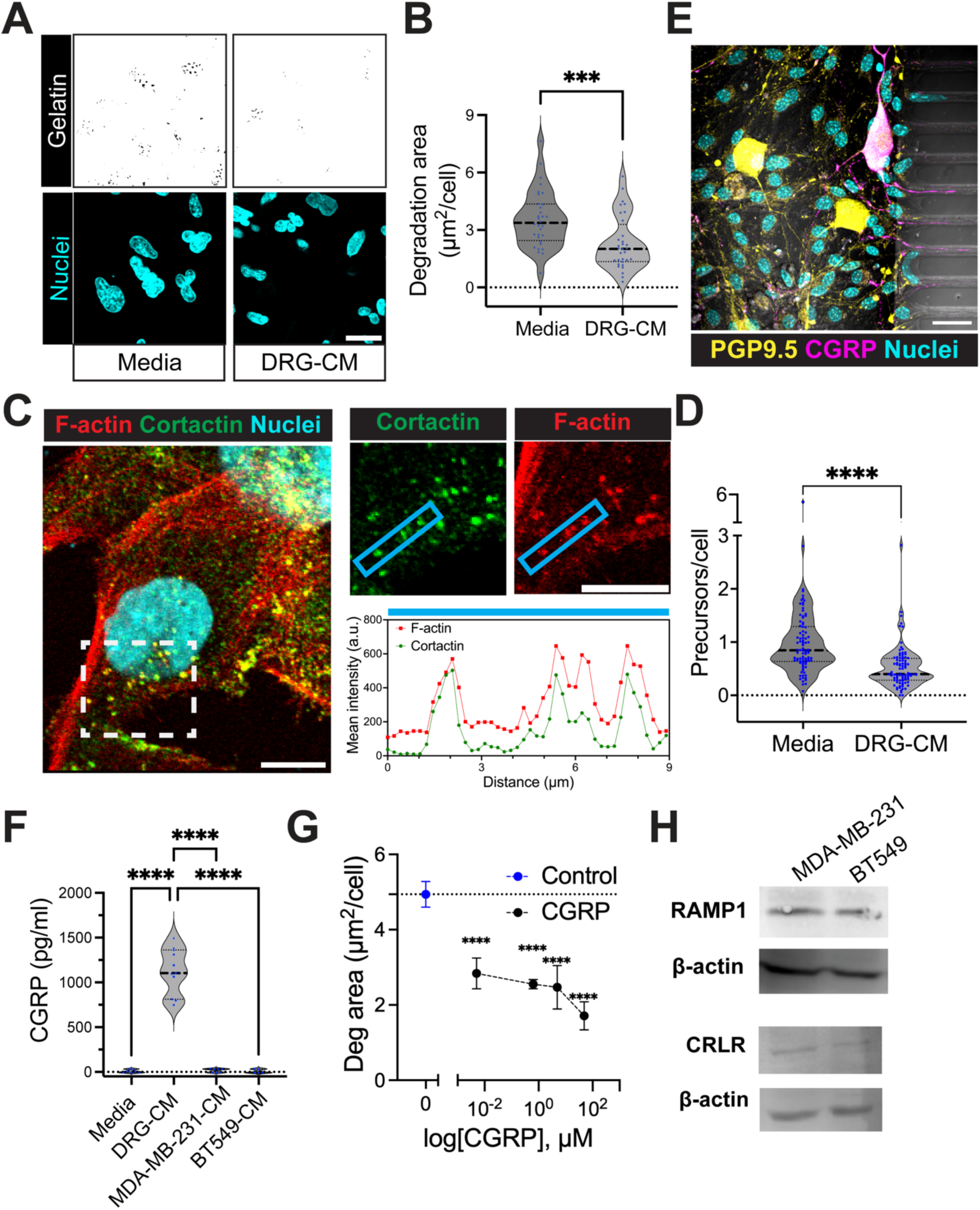
CGRP released by sensory axons inhibits invadopodia assembly and invadopodia-mediated matrix degradation. **A**. Representative images of degraded gelatin by MDA-MB-231 cells cultured in standard medium (left) or DRG-CM. Scale bar = 25 μm. **B**. Quantification of gelatin degradation (μm²/cell) of MDA-MB-231 cells treated with control media or DRG-CM as in A. The graph shows data from three biological replicates (N = 3) with three technical replicates each (n = 3). Statistical significance was determined using a two-tailed Mann-Whitney test (\**p* < 0.05). **C**. Immunofluorescence of MDA-MB-231 cells cultured with DRG-CM. The image shows the nuclei (DAPI, cyan), cortactin (green), and F-actin (phalloidin, red), as well as a zoom-in of the independent channels of Cortactin and F-actin (top right). Scale bar: 10 µm. Bottom right line plot shows the intensity profiles of F-actin (red) and Cortactin (green) signals along the blue square. **D**. Quantification of the number of invadopodia precursors (defined as F-actin and Cortactin colocalizing spots) per cell. The graph shows data from three independent experiments performed in triplicate. Statistical significance was tested with the Mann-Whitney test (\**p* < 0.05). **E**. Immunofluorescence showing the nuclei (DAPI, cyan), neurons (PGP9.5, yellow), and CGRP+ sensory neurons (magenta) on a primary culture of sensory neurons plated on the neuronal compartment of DACIT. **F**. Quantification of Calcitonin gene-related peptide (CGRP) present in DRG-CM, MDA-MB-231 (MDA-MB-231-CM), and BT549-CM cells cultured for 48 hours and control media. Data represent three biological replicates (N = 3, n = 3). Statistical significance was determined using two-way ANOVA, Dunnett’s multiple comparisons test (**** *p* <0.0001). **G**. Quantification of gelatin degradation (μm²/cell) of MDA-MB-231 cells treated with increased concentration of CGRP (5 nM to 50 µM). The graph shows data from three biological replicates (N = 3) with three technical replicates each (n = 3). Statistical significance was determined using Dunnett’s multiple comparison test (\**p* < 0.05). **H**. Western blot analysis of receptor activity-modifying protein 1 (RAMP1) and calcitonin receptor-like receptor (CRLR) expression in human breast cancer cells (MDA-MB-231 and BT549). β-actin was used as a loading control.

We hypothesized that CGRP, a neuropeptide released by the most prevalent sensory neuron subtype in the adult mouse DRG, is responsible for the inhibition of invadopodia assembly and function. To test this, we confirmed the presence of CGRP+ neurons in our DRG primary culture (**Figure 3E**) and verified that primary DRG neurons secreted CGRP in culture, while cancer cells did not (**Figure 3F**). Importantly, the reduction in invadopodia-mediated degradation was dependent on the concentration of CGRP (**Figure 3G**).

The canonical mechanism of CGRP action is ligand-receptor binding, with the principal receptor being a heterodimer composed of receptor activity-modifying protein 1 (RAMP1) and calcitonin receptor-like receptor (CRLR). Western blot analysis (**Figure 3H**) revealed that MDA-MB-231 and BT549 cell lines express both CGRP receptor-related genes: RAMP1 and CRLR, suggesting a direct mechanism by which extracellular CGRP signals to cancer cells.

### CGRP-dependent inhibition of invadopodia is mediated by increase in cAMP

The CGRP receptor complex CRLR-RAMP1 belongs to the large family of G-protein–coupled receptors (GPCRs), which activate G-proteins upon ligand binding. Depending on the cell type and the specific G-protein subunit involved, CRLR-RAMP1 activation can result in distinct signaling pathways, leading to (a) increased cyclic AMP (cAMP), (b) decreased cAMP, (c) elevated intracellular calcium, or (d) enhanced nitric oxide (NO) production ^42^.

We found that Substance P, another sensory neuron–derived neuropeptide whose GPCR signaling increases cAMP ^43^, inhibit invadopodia-mediated gelatin degradation similarly to CGRP (**Supplementary Figure 3**). In contrast, Neuropeptide Y (NPY), which signals through GPCRs reduce cAMP levels ^44^ had no effect on invadopodia mediated degradation (**Supplementary Figure 3**), suggesting that increased cAMP levels may underlie invadopodia inhibition. Kisspeptin-10, a matrikine peptide known to increase invadopodia-mediated gelatin degradation ^45^ was used as a positive control.

To determine whether CGRP modulates invasion via cAMP signaling, we transiently expressed the fluorescent cAMP biosensor, G-Flamp1 ^46^ in MDA-MB-231 cells and monitored fluorescence changes in vehicle-treated cells and following addition of CGRP (**Figure 4A, 4B**). Forskolin, a direct activator of adenylyl cyclase, which increases cAMP levels without the need for preceding increase in CGRP, was used as a positive control. Treatments with either CGRP or forskolin resulted in transient increases in intracellular cAMP values, while cells treated with the vehicle did not induce fluorescence (**Figure 4A, 4B**). Collectively, these data indicated that breast cancer cells respond to CGRP by increasing intracellular cAMP levels.

We next asked whether elevated cAMP levels were sufficient to inhibit invadopodia-mediated matrix degradation as observed with increasing concentrations of CGRP (**Figure 3G**). To test this, we treated cancer cells with forskolin or IBMX, which elevates intracellular cAMP by inhibiting cAMP phosphodiesterase-mediated degradation (**Figure 4C**). Both treatments phenocopied CGRP-mediated inhibition of gelatin degradation, supporting a cAMP-dependent mechanism.

**Figure 4.**
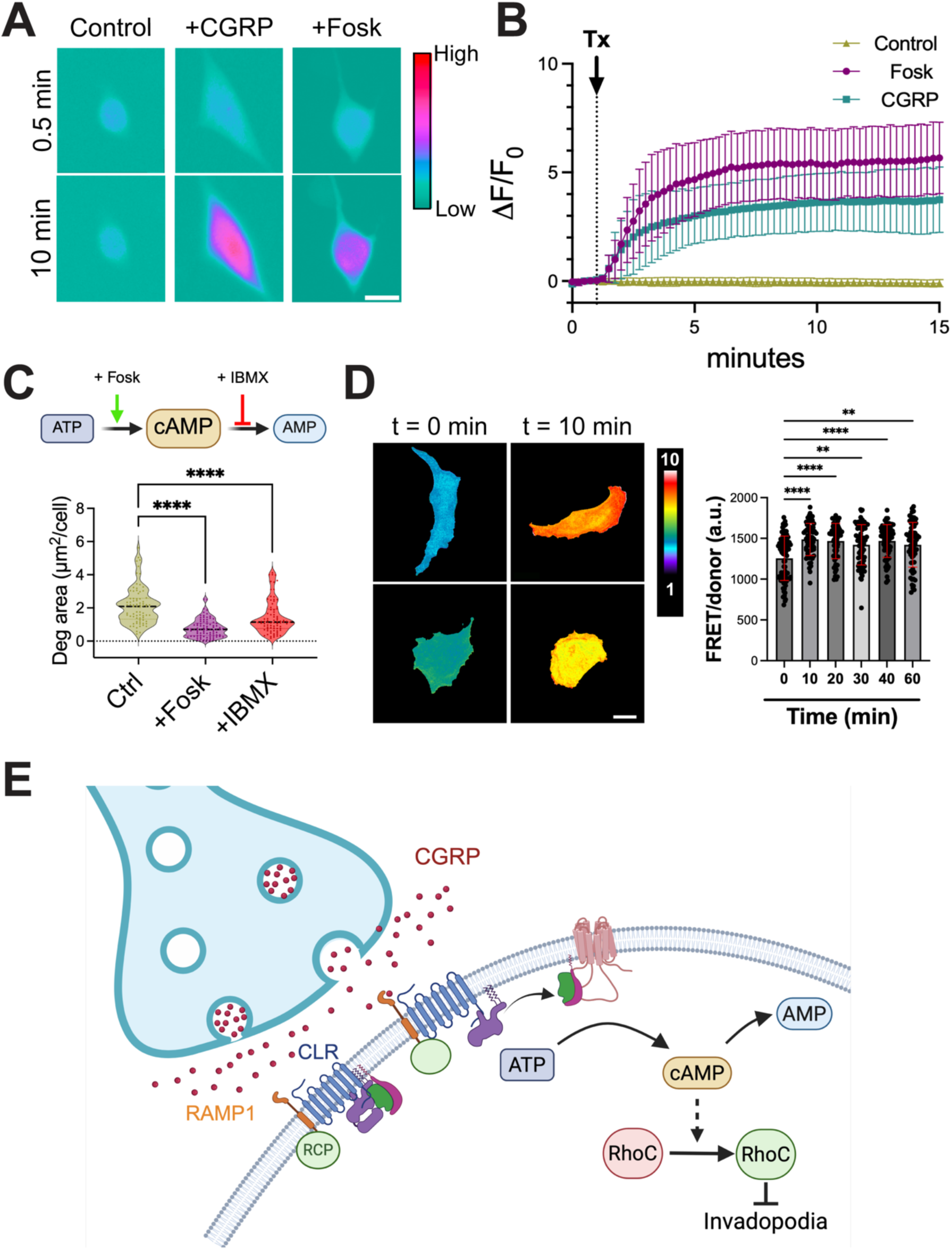
CGRP-dependent inhibition of invadopodia inhibition is mediated by increase in cAMP. **A**. Representative images of MDA-MB-231 cells transiently expressing Flamp1 before (at minute 0.5) and after (at minute 10) stimulation with CGRP (+CGRP, 50 µM) or Forskolin (+Fosk, 40 µM) or vehicle (Control). Scale bar: 25 μm. **B**. Traces of dF/F_0_ Flamp1 protein upon stimulation with 40 µM Forskolin (Fosk, *n*=13) or 50µM CGRP (*n*=17) or vehicle control (*n*=21). Data from three independent biological experiments are presented. **C** Gelatin degradation assay of MDA-MB-231 cells plated with 40 µM Forskolin or 100 µM IBMX (bottom) and scheme (top) showing how Forskolin (Fosk) and IBMX impact gelatin degradation. Outliers were identified with Robust Outlier Removal method and statistical significance was determined using two-way ANOVA, Dunnett’s multiple comparisons test (**** *p* <0.0001). **D**. FRET biosensor analysis of RhoC activity during 40 µM Fosk stimulation, showing activation of RhoC 10 min following the stimulation and thereafter. Representative ratiometric images for T = 0 and T = 10 min time points are also shown, Scale bars = 20 µm. Statistical significance was determined by using two-way ANOVA (**** *p* <0.0001), from aggregate data sets of individual fields of view, compiled from 4 independent experiments, shown with 95% confidence intervals. **E**. Scheme illustrating sensory neuron–secreted CGRP reaches CRLR/RAMP receptors on breast cancer cells and activates adenylate cyclase, leading to increased intracellular cAMP levels resulting in a reduction of invadosome precursors and decreased matrix degradation.

Canonically, cAMP has been linked to Protein kinase A-mediated inactivation of RhoA through phosphorylation at Ser188. This phosphorylation enhances RhoA sequestration by Rho guanine nucleotide dissociation inhibitor (RhoGDI) ^47^. While the effects on RhoA are established, cAMP mediated regulation of RhoC, a close paralog with documented roles in controlling invadopodia in breast cancer cells ^48–50^, remains unexplored. Using an optimized Förster Resonance Energy Transfer (FRET)-based biosensor for RhoC ^20^, we measured the whole-cell activity levels of RhoC in cells under forskolin treatment. In a time-dependent manner, forskolin treatment induced significant whole-cell RhoC activation in MDA-MB-231 cells, which peaked at 10 min and remained elevated throughout the assay (**Figure 4D**). Importantly, these observations clearly distinguish differential signaling landscape surrounding paralog isoforms of Rho GTPase in controlling critical cellular functions and the phenotype.

### Reduced RAMP1 levels promote 3D invasion and metastasis

So far, our data suggest that activation of the CGRP signaling pathway reduces the invasive capacity of cancer cells *in vitro*; however, its effect on breast cancer progression *in vivo* remains unclear. To test the impact of CGRP-mediated signaling pathway on breast tumor metastasis, we generated stable cell lines expressing fluorescent Dendra2, MDA-MB-231-D2, and either *RAMP1* knockdown or shCtrl (**Figure 5A**). Reduced levels of *RAMP1* did not affect the proliferation of MDA-MB-231 cells *in vitro* (**Figure 5B**) but significantly increased the 3D spheroid invasion (**Figure 5C-D**).

**Figure 5.**
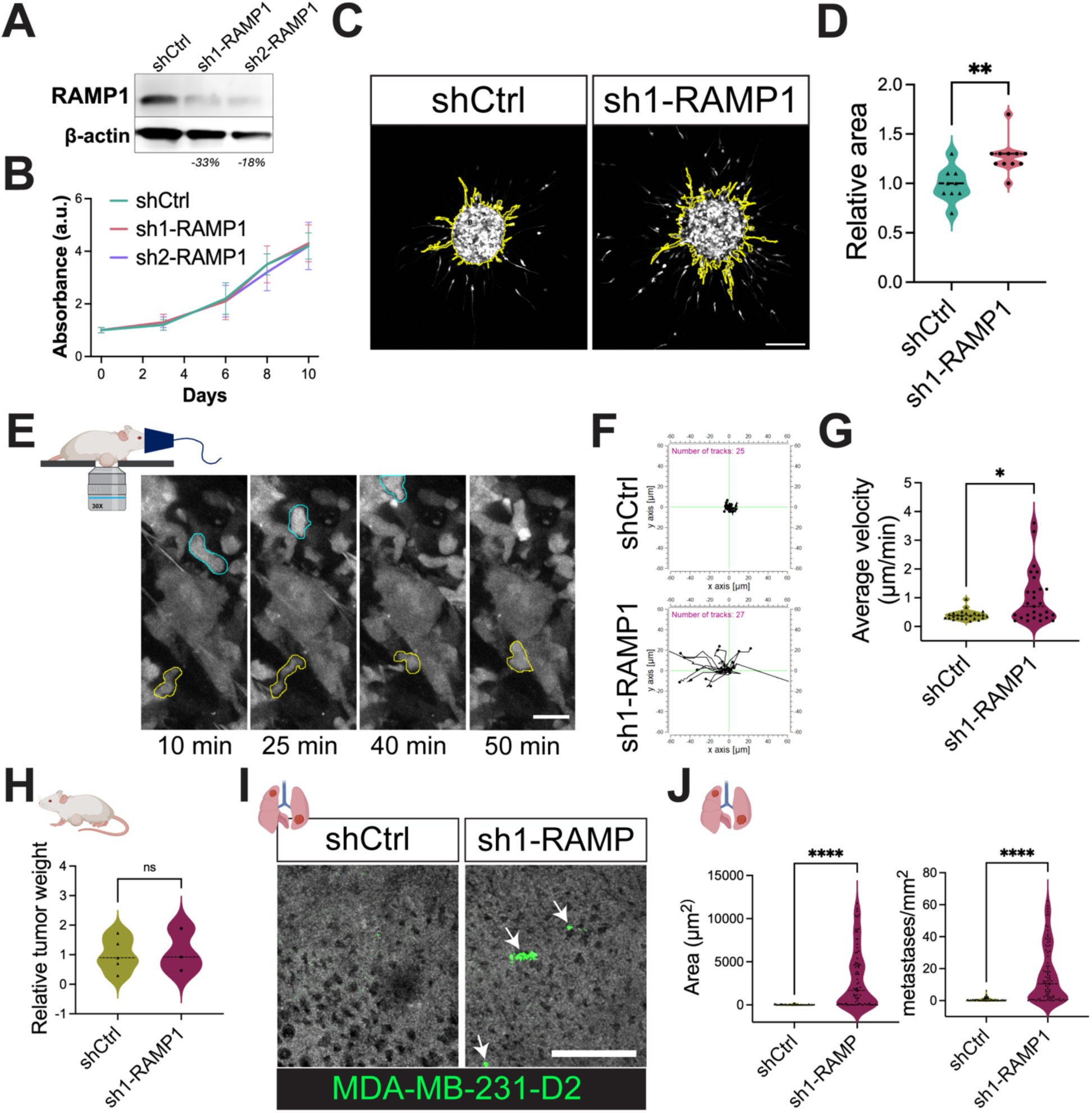
RAMP1 knock down increases 3D spheroid invasion, cell migration and lung metastasis of breast cancer cells. **A**. Western blot of control (shCtrl), sh1-*RAMP1* and sh2-*RAMP1* MDA-MB-231 cells using antibodies against RAMP1, and ß-actin as protein loading control. **B**. Cell proliferation was assessed by crystal violet assay. Graph shows the absorbance units (a.u.) at days 0 (12 h after plating), 3, 6 8, 9 and 10. Data from three biological each with 5 technical replicates (N=3, n=5) are shown. The doubling time (shCtrl: 4.33 days; sh1-*RAMP1*: 4.29 days; and sh2-*RAMP1*: 4.34 days) was calculated using the exponential (Malthusian) growth model. Representative images (**C**) and quantification (**D**) of 3 days invading MDA-MB-231-D2 spheroids. Yellow line shows the invasive mask. Scale bar = 200 µm. **D**. Graph show the relative area of invading spheroids of two biological, ≥ 4 spheroids per group. Statistical differences were determined by Mann-Whitney test. **E.** Representative stills from intravital imaging of MDA-MB-231-D2 sh1-*RAMP1* mammary tumors, showing manually traced migrating cells (blue and yellow masks). Scale bar, 20 µm. Graphs show MDA-MB-231-D2 cells shCtrl (n=25) and sh1-*RAMP1* (n=27) migrating trajectories (**F**) or average speed (**G**, µm/min) of cells during intravital imaging. Statistical differences were determined by Mann-Whitney test. **H**. Graph shows the relative tumor weight determined after dissection. Statistical differences were assessed with the Mann–Whitney test. Representative images (**I**) and quantification (**J**) of lung metastases of control (shCtrl) or sh1-*RAMP1* MDA-MB-231-D2 tumor bearing mice. White arrows point to metastases colonies. Scale bar=200µm. Statistical differences were assessed using the Mann-Whitney statistical test.

To establish orthotopic mouse xenografts, we injected control and *RAMP1* knock-down MDA-MB-231-D2 cells and monitored tumor growth. Using intravital multiphoton microscopy, we then monitored the movement of cancer cells in the areas with intact, flowing blood vessels (**Figure 5E**). Both total distance (**Figure 5F**) and velocity (**Figure 5G**) of sh-RAMP1 cancer cells were increased compared to shCtrl cells, suggesting that reduced levels of RAMP1 increase cancer cell motility *in vivo*.

Following sacrifice (∼70 days, ∼1cm^2^ diameter), tumors and lungs were collected for analysis. Control shCtrl and sh-RAMP1 tumor weights did not show significant differences (**Figure 5H**). However, sh1-RAMP1 cells formed higher number (**Figure 5I, 5J-right)** and larger lung metastases (**Figure 5I, 5J-left**) compared with the control.

Overall, these data suggest that the lack of CGRP receptor signaling favors the invasion of breast cancer cells.

### Invasive breast tumor subtypes have decreased expression of CGRP receptor-related genes in human and mouse

Given that reduced CGRP signaling in cancer cells may promote metastasis, we examined the expression of CGRP receptor components in breast tumors. To distinguish stromal from cancer cell expression, we analyzed publicly available scRNA-seq datasets from mouse (GSE221528) and human (GSE75688) tumors, focusing on genes encoding the CGRP receptor complex: RAMP1 and CALCRL (CRLR).

Both murine and human datasets showed that *Ramp1* and *Calcrl* are expressed across multiple cell types (**Figure 6A, 6B, 6D**), consistent with reports that indicate endothelial cells ^51,52^, fibroblasts ^53^, and immune cells ^54,55^ are sensitive to CGRP. Notably, their expression patterns were discordant. For example, at early stages (**Figure 6A-B day 11**), fibroblasts expressed high levels of *Ramp1* but minimal *Calcrl* whereas cancer cells show the opposite pattern (**Figure 6A-B**), suggesting asynchronous regulation and potentially distinct functional roles.

**Figure 6.**
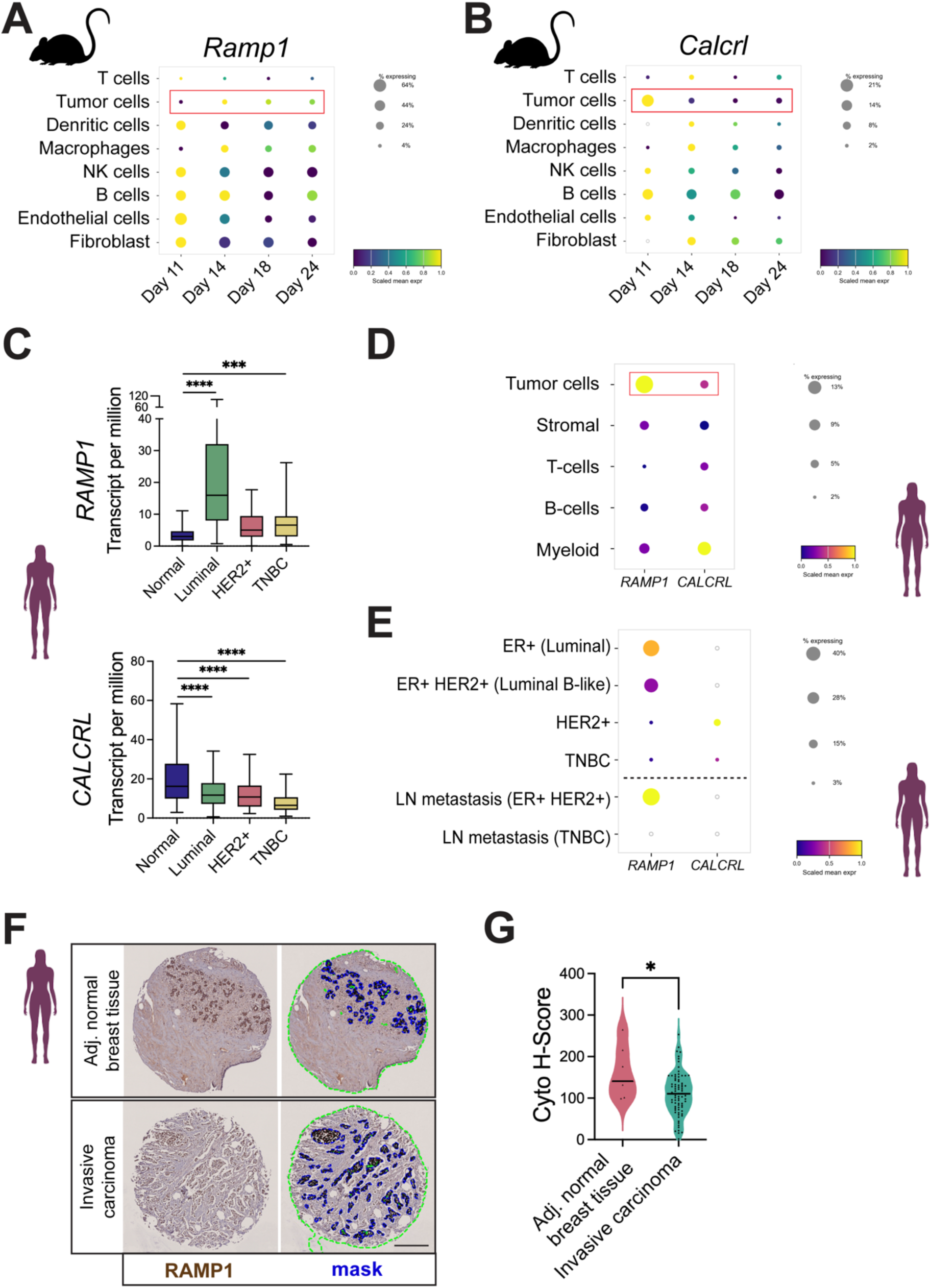
Invasive breast tumor subtypes, in human and mouse, have decreased expression of CGRP receptor-related genes. Dot plots showing expression of *RAMP1* (**A**) and *CALCRL* (**B**) in murine single cell transcriptome data (scRNAseq, GSE221528) at days 11, 14, 18 and 24 after breast tumor cell injection. Data correspond to 2 tumors and 2104 (day 11); 3098 (day 14); 2756 (day 18), 2298 (day 24) cells analyzed. **C**. Expression of *Ramp1* (top) and *Calcrl* (bottom) in human breast tumors from bulk TCGA datasets (UALCAN). Normal (n=114), Luminal (n=566), HER2+ (n=37), TNBC (n=116). Statistical differences were determined by ANOVA/t-tests (**** p <0.0001). **D**. Dot plot showing *Ramp1* and *Calcrl* expression across different cell types in human breast tumors (scRNAseq, GSE75688). **E**. Dot plot showing *Ramp1* and *Calcrl* expression (scRNAseq, GSE75688) in cancer cells from breast cancer subtypes: ER+ (Luminal) (N=2, 73 cells), ER+HER2+ (luminal B-like) (N=1, 15 cells), HER2 (N=3, 130 cells), TNBC (N=4, 63 cells) as well as lymph nodes metastases: ER+HER2+ (N=1, 10 cells), or TNBC (N=1, 26 cells). Dot size represents the fraction of cells with detectable gene expression (> 0), scaled within each plot. Dots representing <1% expression are shown as empty circles. Dot color indicates the mean log-normalized expression within each group. Color scales (min-max) are scaled independently per each cell type (row) and are not comparable across rows or plots. **F**. Representative images and cytoplamic pixel-based H-scores (H-scores) of RAMP1 signal of the epithelial or tumor region of adjacent normal breast tissue (Adj. normal breast tissue, from 7 patients) and invasive carcinoma (from 80 patients) stained with the RAMP1 antibody and counterstained with hematoxylin. Scale bar 300µm. Mann Whitney test was applied to determine statistical significance (* *p*-value = 0.0352).

Longitudinal murine analysis demonstrated that both genes are expressed in tumor cells from early stages (**Figure 6A**). However, *Calcrl* expression and the proportion of *Calcrl*-positive tumor cells decreased from day 11 to day 24, whereas *Ramp1* expression levels fluctuates (**Figure 6B**).

Human bulk RNA-seq analysis revealed tumor subtype-dependent variation in *RAMP1* and *CALCRL* expression, with the lowest levels observed in aggressive subtypes (HER2+ and triple-negative breast cancer, TNBC) compared with less aggressive tumors (Luminal) (**Figure 6C**). Importantly, human single-cell resolved datasets showed that *RAMP1* was predominantly expressed in cancer cells, whereas *CALCRL* was enriched in myeloid cells (**Figure 6D**).

Subtype comparison further showed that less aggressive tumors (Luminal and Luminal B-like) contained a higher proportion of *RAMP1*-expressing cancer cells with elevated expression levels compared to HER2+ and TNBC tumors (**Figure 6E**), suggesting that *RAMP1* downregulation may contribute to tumor aggressiveness.

To validate this observation at the protein level, we quantified RAMP1 expression within epithelial regions of invasive carcinoma (n=80) and adjacent normal breast tissue (n=7) in patient TMAs (**Figure 6F, masked area**). Cyto H-score quantification revealed that RAMP1 expression was significantly reduced in invasive carcinoma relative to adjacent normal tissue (**Figure 6G**).

Collectively, our findings suggest that aggressive breast tumor cells progressively lose RAMP1 expression, potentially as a mechanism of evasion of neuronal inhibition.

## Discussion

The involvement of peripheral nerves in tumor growth and progression has gained increased attention in recent years. Although recent studies have established the presence and relevance of neurons within breast tumors, relatively few studies have investigated the roles of sensory neurons, and their impact on breast cancer cells remains unclear. While some reports suggest that sensory neurons suppress breast tumor growth ^4,5^, others suggest that they may promote breast tumor progression and metastasis ^7,53,56^. These apparent discrepancies may reflect the multifaceted roles of sensory neurons, which can influence not only cancer cell behavior but also various stromal components, including fibroblasts, immune and endothelial cells.

Here, we show that murine mammary tumors recruit CGRP+ sensory neurons, a subset of which localize close to vessels, where they are known to regulate blood pressure and vascular function ^57,58^. Previously, we showed that invadopodia assemble in contact with the perivascular matrix^18^. Given this spatial overlap, we investigated whether sensory neurons regulate invadopodia-mediated matrix degradation, a key step that precedes intravasation and metastasis ^18,59^. By isolating sensory axon–cancer cell interactions, we found that sensory neuron-secreted CGRP inhibits invadopodia assembly and matrix degradation in triple-negative breast cancer cells via upregulation of cAMP.

Invadopodia inhibition was more pronounced in the presence of axons, compared to the DRG-conditioned medium alone, suggesting that the presence of cancer cells may be increasing the secretion of CGRP. While we observed sensory neurons secrete CGRP at detectable levels in the absence of cancer cells, a few previous studies reported that interactions with cancer cells may be enhancing sensory neuron activities. For example, cancer cells were reported to increase neuropeptide secretion in the tumor-bearing mice ^60^, and increase sensitization upon co-culture with melanoma cells^61^ and breast cancer cells ^53^.

The relative decrease in degradation levels corresponds to local concentration of 100 µM in cancer cells contacted by axons, and only 10 nM in cancer cells cultured in DRG-CM, based on our CGRP dose response measurements. As we have not identified a correlation between axonal distance and the number or size of degradation puncta, such disparity may suggest that the sustained source of CGRP, provided by axonal contacts, replenishes degraded CGRP, as neuropeptides typically have a short half-life in culture ^62^.

Mechanistically, we show that sensory neuron-secreted CGRP increases intracellular cAMP in breast cancer cells. Since it has been reported that other neuropeptides as Substance P, also increase cAMP intracellular levels ^43^, this mechanism may be a more general pattern of neuropeptide effect. Increased cAMP levels were previously shown to reduce migration of breast cancer cells^63^. Here, we demonstrate that increased intracellular cAMP suppresses invadopodia-mediated degradation. Our data also show that DRG-conditioned medium reduced invadopodia assembly, collectively suggesting that CGRP impairs invadopodia assembly and function by inhibiting actin polymerization.

Previous works revealed that increased cAMP reduces RhoA activity through Protein Kinase A (PKA) phosphorylation ^64^. PKA-mediated phosphorylation has been shown to directly modulate the binding affinity of guanine nucleotide dissociation inhibitor (GDI) toward RhoA, accelerating its extraction from the plasma membrane into the cytosol ^65^. Here, we observe that elevated cAMP levels activate RhoC, which is unexpected. However, subtle differences within the switch I and the insert regions of RhoA versus RhoC may confer a preferential affinity of GDI for RhoA over RhoC, under PKA-phosphorylated states. Such differential binding could bias the system toward RhoA inhibition while allowing RhoC activation, given that cellular GDI is typically expressed at approximately equimolar concentrations relative to its target Rho GTPases under basal conditions ^66,67^. Because RhoC expression levels are significantly elevated in migratory and metastatic breast tumor cells ^68^ preferential binding and extraction of RhoA by GDI could impact the local stoichiometric balance of available GDI. This redistribution would reduce effective GDI buffering of RhoC, thereby promoting increased RhoC activation under conditions of elevated cAMP.

Spatiotemporal RhoC activation regulates invadopodia by inactivating cofilin in the surrounding ring, thereby confining active cofilin to the invadopodium core. This leads to enhanced structural integrity and enabling controlled matrix degradation^50^. In addition, activated RhoC establishes an asymmetric zone that restricts cofilin-mediated actin barbed-end formation to a narrow band at the leading edge, promoting efficient cell motility^49^. In contrast, RhoA forms a thin zone of activation at the extreme tip of the leading edge, where it likely engages distinct actin regulators, including formins, and establishes contractile polarity through activation of actomyosin pathways ^69^. Importantly, previous studies have shown that the balance between these spatially segregated RhoA and RhoC activity zones is critical for controlling breast cancer cell motility and invasion ^49,70^. The mechanism observed here appears to disrupt this balance by acutely activating RhoC in response to elevated cAMP, while concomitantly inhibiting canonical RhoA signaling. Such conditions would shift the equilibrium of actin regulation toward the RhoC–ROCK–LIMK–cofilin pathway and away from RhoA-mediated actomyosin contractility, thereby impairing the coordinated integration of invasive, protrusive, and contractile processes required for efficient migration and invasion. Future studies employing spatially resolved RhoC biosensors in the presence of cAMP agonists could directly test whether CGRP induced cAMP elevation disrupts the characteristic ring pattern of RhoC activation at invadopodia.

In the scRNAseq data, components of the CGRP receptor complex, RAMP1 and CRLR, do not shown synchronized expression. This could be explained by each of the components also being involved in interactions with other proteins, or activating alternate pathways. For example, CRLR can interact with RAMP2 or RAMP3, forming the adrenomedullin receptor^71,72^, while RAMP1 and CRLR can also interact a functional amylin receptor^73^.

Interestingly, while cancer cells downregulate *Ramp1* and *Calcrl* expression during tumor progression, both MDA-MB-231 and BT549, representing metastatic human triple negative cells, express a certain level of functional CGRP receptors. This suggests that a low percentage of aggressive breast cancer retains a level of responsiveness to CGRP-mediated signaling. Alternatively, as not all the blood vessels in tumors are innervated by CGRP+ axons, cancer cells sensitive to CGRP may be able to intravasate into CGRP-blood vessels.

Our scRNA analyses revealed that *Ramp1* and *Calcrl* are expressed in stromal cells at similar levels and frequency than in cancer cells. In fact, RAMP1 was proposed as a bad prognosis marker in osteosarcoma, correlating with higher infiltration of gamma-delta T cells ^74^.

Finally, single-cell analysis demonstrated that both CRLR and RAMP1 are highly expressed in stromal cells. In line with this, several studies have reported that CGRP, secreted by sensory neurons, influences stromal cell behavior and contributes to increased tumor aggressiveness. For instance, Zhang et al. found that CGRP stimulates breast fibroblasts, resulting in higher matrix density and promoting immune exclusion ^53^. Additionally, other studies have shown that CGRP suppresses T cell activity in melanoma ^61^ and head and neck squamous cell carcinoma ^75^. In fact, the systemic inhibition of the CGRP-signaling pathway with the CGRP antagonist Rimegepant reduced tumor volume and increased the survival in murine gastric tumor models^76^. These findings indicate that therapeutic strategies should account for the fact that, in addition to cancer cells, tumor stromal cells, such as immune, endothelial cells, and fibroblasts also respond to neuropeptides secreted by activated sensory neurons, thereby influencing tumor progression in a broader context ^61,75^.

In addition to CGRP and substance P, other cAMP activators may also impact cancer cell invasion. Given that multiple GPCRs can modulate cAMP levels, it is worth exploring whether alternative pathways could mimic or enhance the effects observed with CGRP. Accordingly, some reports proposed forskolin as an anti-tumorigenic therapy ^77^. Examining how sensory neuron–stromal cell interactions contribute to cancer cell behavior and tumor progression will likely identify new targets for metastatic breast cancer and yield therapeutic benefits.

## Author contributions

Conceptualization, I.V.-Q. and B.G.; sample preparation, I.V.-Q., E.B., A.J., D.B.V.C., R.F., M.C.M.C., Y.D.; data acquisition, analysis, and interpretation, I.V.-Q., E.B., E.C., A.J., and B.G.; writing, I.V.-Q., E.B., L.H. and B.G.; supervision, E.C., L.H. and B.G.

## Acknowledgements

We thank Kathy Q. Cai, MD, PhD. In the Histopathology Facility of Fox Chase Cancer Center for her help in pathological assessment of patient tissues.

We thank the following funding sources for their support: NIH NCI R01 CA230777; American Cancer Society Research Scholar grant 134415-RSG-20-34-01-CSM, DOD BCRP Breakthrough Award BC230197, PA-CURE, W. W. Smith Charitable Trust awards and STEM Research Award 2025 to B.G., METAvivor Metastatic Breast Cancer Early Career Research Award to I.V.Q., and R35GM132662 from NIH NIGMS to L.H.

---

**Supplementary Figure 1.**
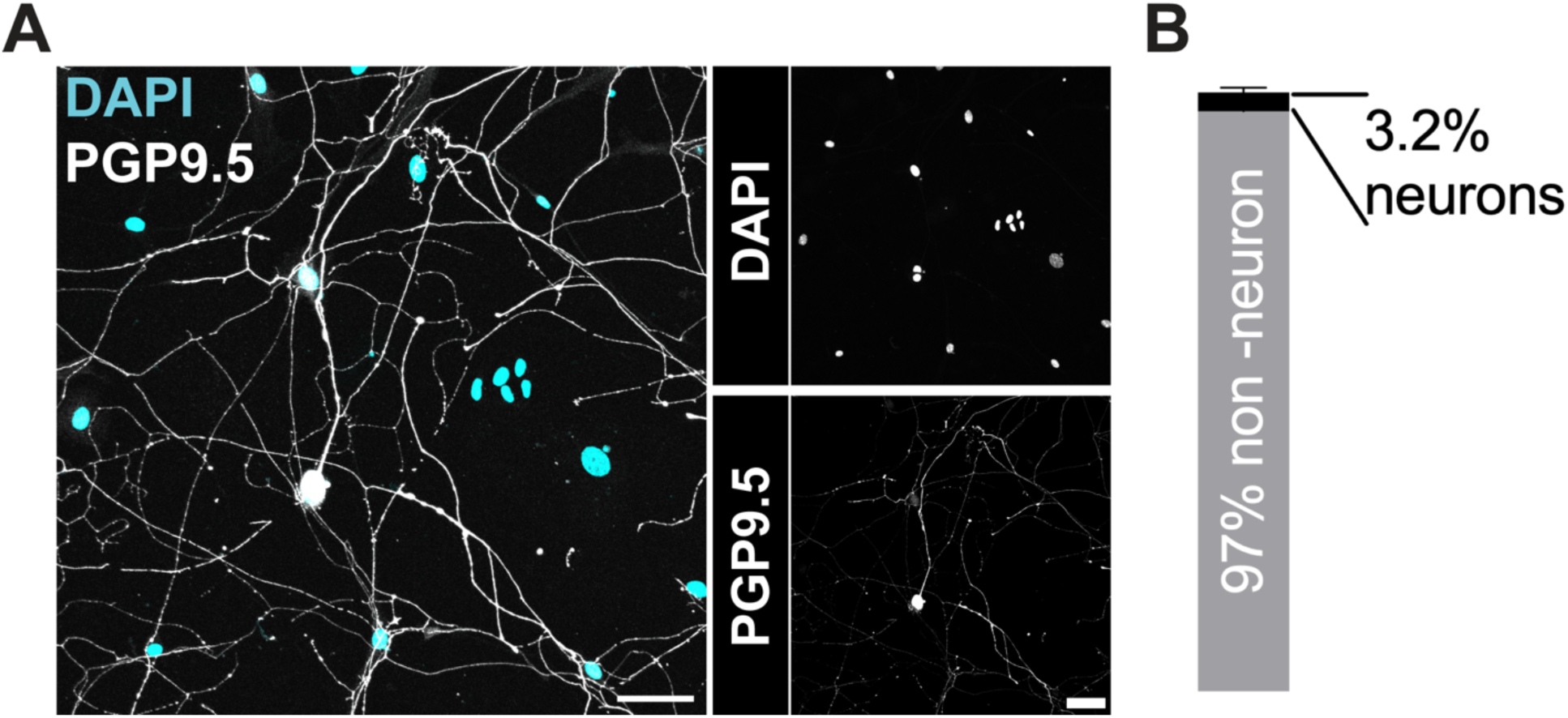
Sensory neurons are a minority subpopulation in the DRG primary cultures. Representative images (**A**) and quantification (**B**) of the percentage of neurons (PGP9.5+) and non-neural cells (DAPI+, PGP9.5-) in a primary culture of DRG cells plated on Matrigel (1:10) for 5 days. Scale bar: 50 µm. The graph shows the percentage of neurons (mean and SD; N=3, n=25) determined as the percentage of PGP9.5+ cells over the total number of cells (DAPI+) per field-of-view.

**Supplementary Figure 2.**
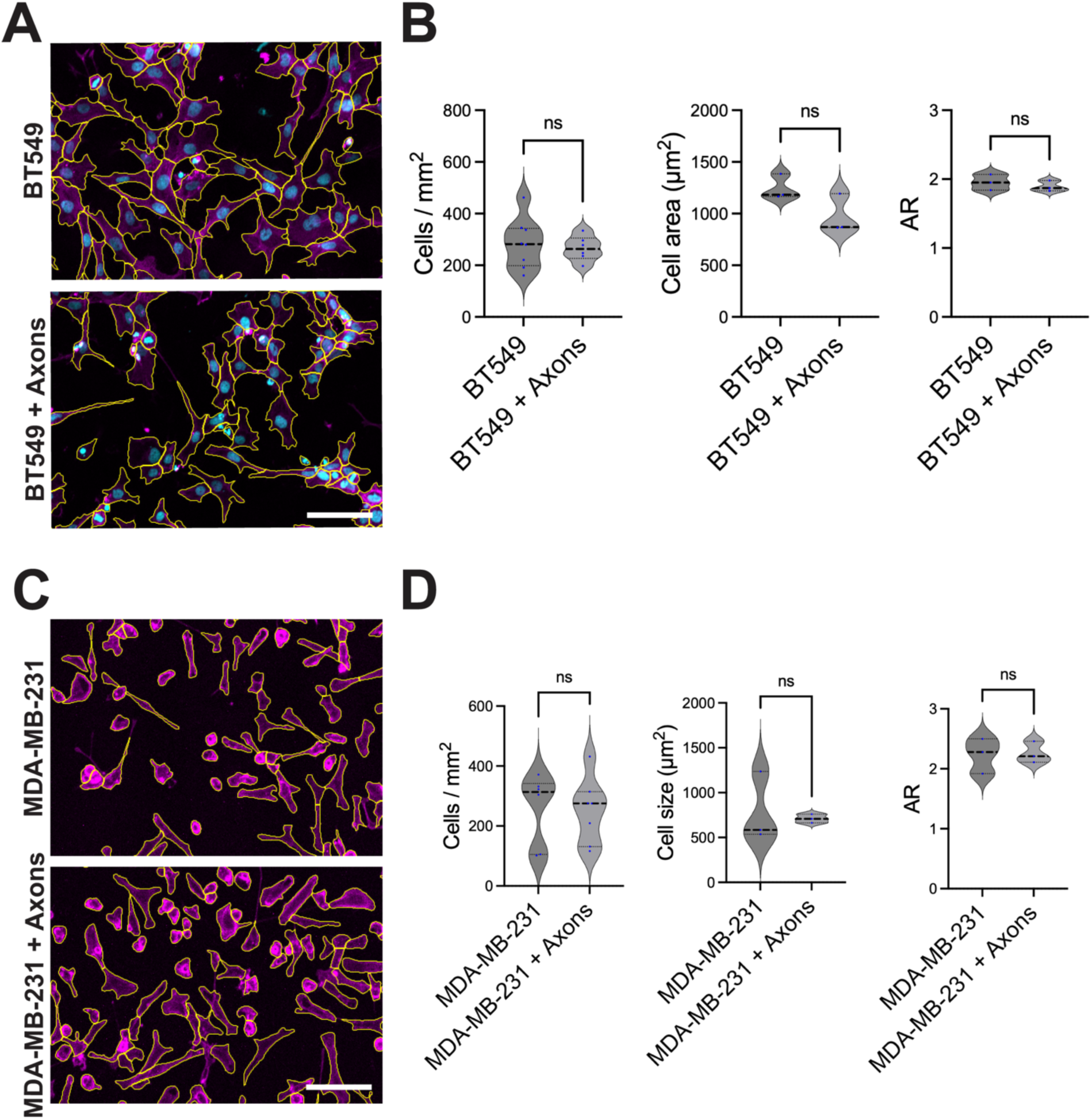
Sensory axons do not affect the proliferation or morphology of breast cancer cells. Representative images (**A** and **C**) and quantification (**B** and **D**) of the number of cells/ mm^2^, the size of the cells (µm^2^) and the aspect ratio (AR) of the BT549 (top) and MDA-MB-231 (bottom) cells with or without axons on DACIT. Scale bar: 100 µm

**Supplementary Figure 3.**
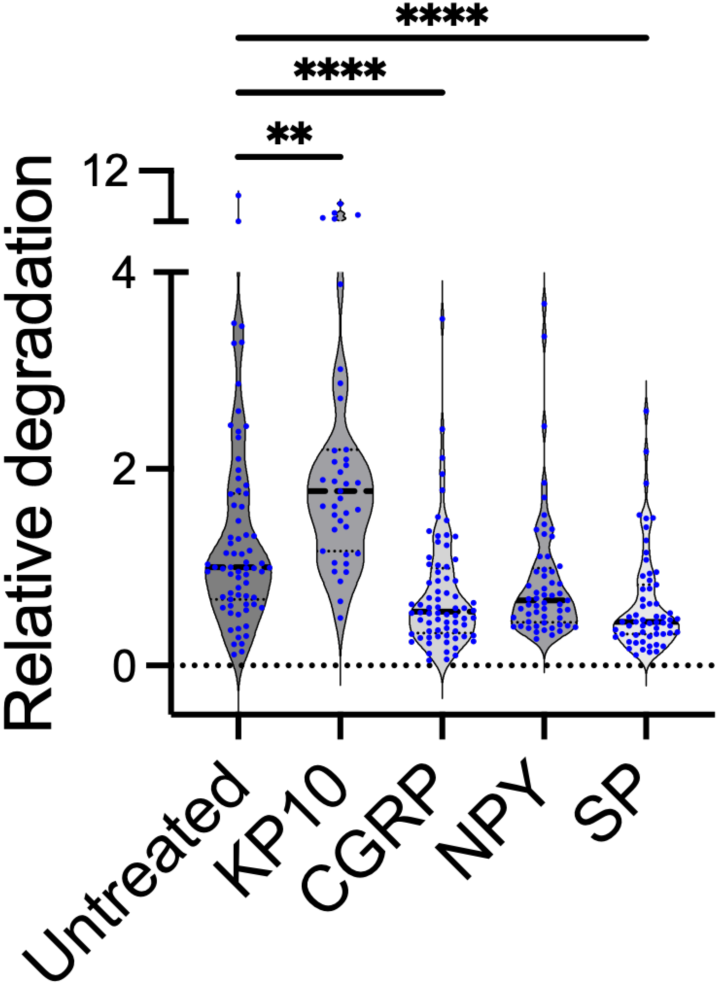
Neuropeptides secreted by sensory neurons decrease invadopodia-mediated matrix degradation. The graph shows the relative gelatin degradation of MDA-MB-231 cells treated with neuropeptides: CGRP, Neuropeptide Y (NPY) or Substance P (SP), all at 5 μM. A matrikine peptide known to promote ECM degradation, Kisspeptin-10, was used as a positive control. Data represent three biological replicates (N = 3, n = 3). Statistical analysis was performed using a two-tailed Mann-Whitney test (*P* < 0.05) comparing each peptide with the untreated cells.

